# Effects of physiological parameter evolution on the dynamics of tonic-clonic seizures

**DOI:** 10.1101/821330

**Authors:** Farah Deeba, Paula Sanz-Leon, P. A. Robinson

**Affiliations:** School of Physics, University of Sydney, NSW 2006, Australia; Center for Integrative Brain Function, University of Sydney, NSW 2006, Australia; Department of Physics, Dhaka University of Engineering and Technology, Gazipur 1700, Bangladesh

## Abstract

The temporal and spectral characteristics of tonic-clonic seizures are investigated using a neural field model of the corticothalamic system in the presence of a temporally varying connection strength between the cerebral cortex and thalamus. Increasing connection strength drives the system into ∼ 10 Hz seizure oscillations once a threshold is passed and a subcritical Hopf bifurcation occurs. In this study, the spectral and temporal characteristics of tonic-clonic seizures are explored as functions of the relevant properties of physiological connection strengths, such as maximum strength, time above threshold, and the ramp rate at which the strength increases or decreases. Analysis shows that the seizure onset time decreases with the maximum connection strength and time above threshold, but increases with the ramp rate. Seizure duration and offset time increase with maximum connection strength, time above threshold, and rate of change. Spectral analysis reveals that the power of nonlinear harmonics and the duration of the oscillations increase as the maximum connection strength and the time above threshold increase. A secondary limit cycle at ∼ 18 Hz, termed a saddle-cycle, is also seen during seizure onset and becomes more prominent and robust with increasing ramp rate. If the time above the threshold is too small, the system does not reach the 10 Hz limit cycle, and only exhibits 18 Hz saddle-cycle oscillations. It is also seen that the times to reach the saturated large amplitude limit-cycle seizure oscillation from both the instability threshold and from the end of the saddle-cycle oscillations are inversely proportional to the square root of the ramp rate.

**Author Summary:** Epilepsy, which is characterized by recurrent seizures, affects around 1% of the world population at some point in their lives. Tonic-clonic seizures are the most commonly encountered primary generalized seizures and it is widely considered that they can be induced by an increase in the connection strength between the cerebral cortex and the thalamus. In this paper, we analyze the detailed dynamics of tonic-clonic seizures along with their dependence on the parameters of the changing connection strength. We study the relationship of the seizure onset, offset, oscillation strength, and oscillation frequency to the duration, amplitude, and rate of change of the connection strength. A detailed understanding of the dynamics and their dependence on the physiological parameters of the brain may explain the variability of seizure dynamics among patients. It may also help to constitute successful seizure prediction.

## Introduction

Tonic-clonic seizures, formerly known as grand mal seizures, are the most frequently encountered generalized seizures [1]. These seizures have a tonic phase, which is characterized by an initial increase in tone of certain muscles, followed by a clonic phase, which involves bilateral symmetric jerking of the extremities [2]. Tonic-clonic seizures have markedly different pre-and post-ictal electroencephalograms (EEG) and typically last 1 to 3 minutes. Primary generalized seizures, which is one of the most commonly seen seizures, begin simultaneously across the whole cortex [1].

A number of authors have investigated the mechanisms of seizures using the neural network and neural field approaches [3–13]. Many authors have proposed that transitions from healthy state to the seizure state occur via bifurcations upon changing physiological parameters [3–9, 12, 13]. For example, depending on the instability region, increasing excitatory connection strengths between cortex and thalamus drives the system into ∼ 10 Hz and ∼ 3 Hz seizure oscillations via a subcritical and supercritical Hopf bifurcation, respectively, once a critical value (i.e., a threshold) is passed [3–9, 12, 13]. Results from *in vivo* studies have provided evidence that changes in corticothalamic connection strengths can induce seizures [12, 14–16], which possibly occur due to a key cellular event triggered by GABA_*B*_ (metabotropic transmembrane receptors for gamma-aminobutyric acid) mediated mechanisms underlying the reduction of the threshold for Ca^2+^ spikes [1, 2] due to the effects of drugs, excess or deficiency of neurotransmitters or neuromodulators [1, 2, 17]. However, the detailed dynamics of generalized tonic-clonic seizure including its dependence to the changing profile of the corticothalamic connection strength have never been studied in detail. The dependence of the spectral characteristics like the frequencies of the oscillations on the parameters of the changing connection strength have also not been studied.

In this study, we apply a widely used neural field model of the corticothalamic system to study the dynamics of tonic-clonic seizures [3–5, 7, 8, 18–20]. Neural field theory (NFT) is a continuum approach that predicts the average dynamics of large numbers of neurons [21, 22]. The specific model used here [23–26] has reproduced and unified many observed features of brain activity based on the physiology, including evoked response potentials [27], activity spectra [28], arousal state dynamics, age-related changes in the physiology of the brain [29], and many other phenomena [3–5, 7, 8, 18–20, 30–32]. The above NFT model has also been used in seizure studies [3–5, 7], where it has successfully unified features of tonic-clonic and absence seizures [3–5, 7], and explain the dependence of the dynamics and interictal oscillations during absence seizures on the parameters of the changing connection strength between the cortex and the thalamus [33, 34]. Previous studies have shown that a gradual increase of the connection strength between the cortex and thalamus near the alpha instability boundary shown in [8] in this model can initiate nonlinear dynamics whose characteristics closely resemble those of tonic-clonic seizures as a result of a subcritical Hopf bifurcation that destabilizes the ∼ 10 Hz alpha resonance [3, 4, 19, 31]. Changes in other connection strengths also introduce similar dynamics because of the universality properties of the Hopf bifurcation [12].

The general property and bifurcation mechanism of the resultant tonic-clonic seizure has been studied in detail in [3]. However, the impact of underlying parameter changes of the corticothalamic connectivity strength on tonic-clonic seizure onset, dynamics, and termination have not been studied in detail. In particular, an extensive study like [33] on the dependence of the onset and termination of tonic-clonic seizure on the temporal form of the connection strength is necessary to understand the variability in seizure events, such as difference in the onset time and duration among different subjects, and to help lay the foundations for tonic-clonic seizure control strategies. These analysis are also necessary to explain the changes in harmonic structures seen in previous studies [35–37] during seizure. In sort, the aims are to understand the effects of physiological parameters on the temporal and spectral characteristics of seizure dynamics, including saddle-cycle oscillations [19].

The outline of this paper is as follows: In the Results, we explore the general characteristics of seizure as well as the dependence of seizure dynamics on the temporal variation of connection strength. In the Discussion, we provide a summary and discuss possible applications of our outcomes and finally, in the Methods section, we present the corticothalamic neural field model along with the temporal variation function and the numerical methods.

## Results

In this section we investigate the dynamical characteristics of model tonic-clonic seizures as well as the effects of the temporal variation of the corticothalamic connection strength, *ν*_*se*_ on the dynamics. For the investigation of general characteristics, we keep a constant maximum connection strength *ν*_max_, characteristic duration *t*_2_ − *t*_1_, and characteristic rise time Δ, and all other parameters listed in Table 1.

**Table 1.** Nominal parameters of the neural field model from [3].

| Parameter | Value | Unit | Meaning |
| --- | --- | --- | --- |
| $\nu_{ee}$ | 1.2 | mV s | Excitatory corticocortical connectivity |
| $\nu_{ei}$ | -1.8 | mV s | Inhibitory corticocortical connectivity |
| $\nu_{es}$ | 1.4 | mV s | Specific thalamic to cortical connectivity |
| $\nu_{re}$ | 0.2 | mV s | Cortical to thalamic reticular connectivity |
| $\nu_{rs}$ | 0.2 | mV s | Specific to reticular thalamic connectivity |
| $\nu_{se}$ | 1.0 | mV s | Cortical to specific thalamic connectivity |
| $\nu_{sr}$ | -1.0 | mV s | Reticular to specific thalamic connectivity |
| $\nu_{sn}\phi_n$ | 2.0 | mV | Subthalamic input |
| $Q_{\max}$ | 250 | s<sup>-1</sup> | Maximum firing rate |
| $\theta$ | 15 | mV | Mean neuronal threshold |
| $\sigma$ | 6 | mV | Threshold standard deviation |
| $\gamma_e$ | 100 | s<sup>-1</sup> | Damping rate |
| $\alpha$ | 60 | s<sup>-1</sup> | Decay rate of membrane potential |
| $\beta$ | 240 | s<sup>-1</sup> | Rise rate of membrane potential |
| $t_0$ | 80 | ms | Corticothalamic return time (complete loop) |
| $t_1$ | 100 | s | Center of the ramp rise |
| $t_2$ | 200 | s | Center of the ramp fall |
| $\nu_{\max}$ | 1.2 | mV s | Maximum value of $\nu_{se}$ |
| $\nu_0$ | 0.8 | mV s | Minimum value of $\nu_{se}$ |
| $\Delta$ | 10 | s | Characteristic rise time |

To investigate the effect of the variation of *ν*_*se*_ on seizure dynamics we vary *ν*_max_, Δ, and *t*_2_ − *t*_1_ individually by keeping all other parameters constant. Figure 1(a) shows the variation of *ν*_*se*_ with time for the parameters values specified in Table 1.

**Fig 1.**
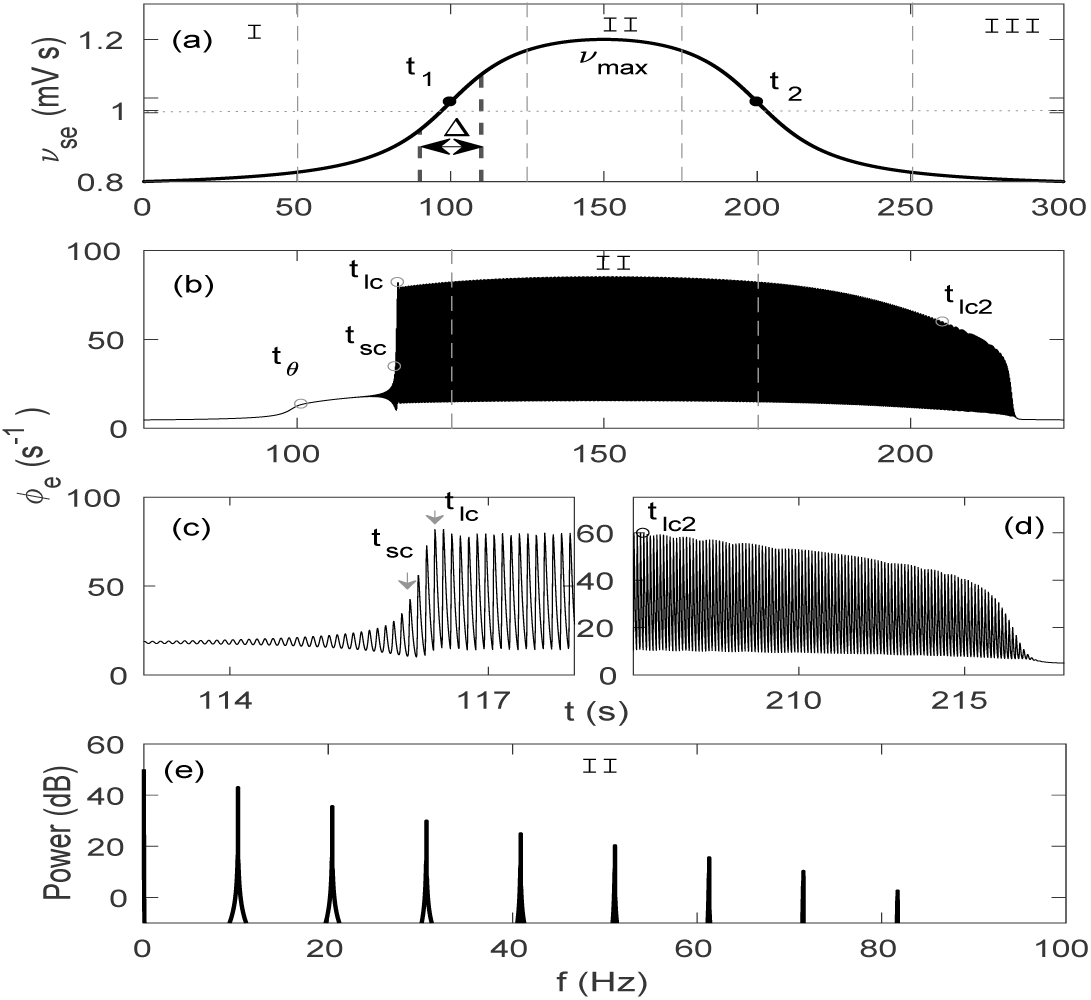
Corticothalamic dynamics for temporally varying *ν*_*se*_, with Δ = 20 s and rest of the parameters shown in Table 1. (a) Temporal profile of *ν*_*se*_ varying from *ν*_0_ to *ν*_max_ and back. Three different regions are identified as: I = pre-ictal state, II = ictal state, and III = post-ictal state. (b) Cortical excitatory field *ϕ*_*e*_ vs. *t*, showing a 10 Hz spike-wave oscillation. Individual oscillations can not be distinguished on this scale. (c) Zoom of *ϕ*_*e*_ at seizure onset. (d) Zoom of *ϕ*_*e*_ at seizure offset. (e) Power spectrum of *ϕ*_*e*_ in Region II. An arbitrary dB scaling is used because clinical EEG recordings involve additional attenuation by structures between the cortex and the electrode, which we do not model here.

### General characteristics of tonic-clonic seizures

Three main regions are distinguished according to the dynamics of the cortical activity *ϕ*_*e*_ (cortical excitatory field) as illustrated in Fig. 1(b): Region I from 0 − 50 s is the pre-ictal state when *ν*_*se*_ is too small to initiate seizure-like oscillations; Region II from 125 − 175 s is the ictal state when *ν*_*se*_ is around its maximum value, *ν*_max_, and the system oscillates with maximum amplitude; and Region III from 250 − 330 s is the post-ictal state, where *ν*_*se*_ returns to its baseline value, and oscillations start decreasing in amplitude until they completely cease.

Figures 1(c) and (d) show the zoomed seizure onset and offset, respectively, which are the transitions from Region I to II, and from Region II to III, respectively.

The normalized power spectrum in Region II is shown in Fig. 1(e). Figure 1(e) shows a dominant resonance at ∼ 10 Hz with multiple harmonics in Region II, where power decreases gradually with frequency.

#### Dynamics of seizure onset

Figure 1(b) shows that in Region I, the system remains in the steady state because *ν*_*se*_ is below the bifurcation threshold. A small increase in *ϕ*_*e*_ due to the increase of *ν*_*se*_ is also seen in this region. At *t* = *t*_*θ*_, which is the time at which *ν*_*se*_ crosses the linear instability threshold, the fixed point loses its stability, and ∼ 18 Hz oscillations appear. The first few oscillations are too small to be distinguished on this scale, but their envelope increases exponentially until *t* = *t*_*sc*_, when the trajectory spirals further outwards to a large amplitude 10 Hz limit cycle, as seen in Fig. 1(c); these 18 Hz oscillations are termed saddle-cycle oscillations because they are due to a saddle cycle located between the stable steady state and the stable large amplitude limit cycle attractor. The envelope of the 10 Hz oscillations continues to increase from *t* = *t*_*sc*_ until *t* = *t*_*lc*_, when the system reaches the large amplitude limit cycle. At *t* ≈ *t*_*lc*_, the amplitude of the oscillations overshoots because *ν*_*se*_ is still rapidly increasing. Then, the amplitude of the oscillations increases gradually until *ν*_*se*_ = *ν*_max_ in Region II, then decreases.

Figure 1(c) shows a clearer view of saddle-cycle oscillations, and times *t*_*sc*_ and *t*_*lc*_; where we define *t*_*lc*_ to be the point of inflection.

#### Dynamics of seizure offset

In Fig. 1(d), we see that the amplitude of the oscillations decreases gradually from its peak during the ramp down of *ν*_*se*_. More specifically, at *t* = *t*_*lc*2_, when *ν*_*se*_ crosses the offset bifurcation threshold *ν*_*lc*2_ = 0.98 mV s [3], the large limit cycle loses stability and the oscillation amplitude decreases steeply to approach the stable steady state in Region III.

#### Differences between onset and offset dynamics

Comparing Fig. 1(c) with Fig. 1(d), we see that *ν*_*θ*_ *> ν*_*lc*2_, as expected for transitions due to a subcritical Hopf bifurcation. This is further seen in Fig. 2, where we see that the system bifurcates from the fixed point at *ν*_*se*_ = *ν*_*θ*_ and reaches the saturated large amplitude attractor at *ν*_*se*_ = *ν*_*lc*_. As *ν*_*se*_ decreases, the large amplitude attractor becomes unstable at *ν*_*se*_ = *ν*_*lc*2_ and the system returns toward the fixed point.

**Fig 2.**
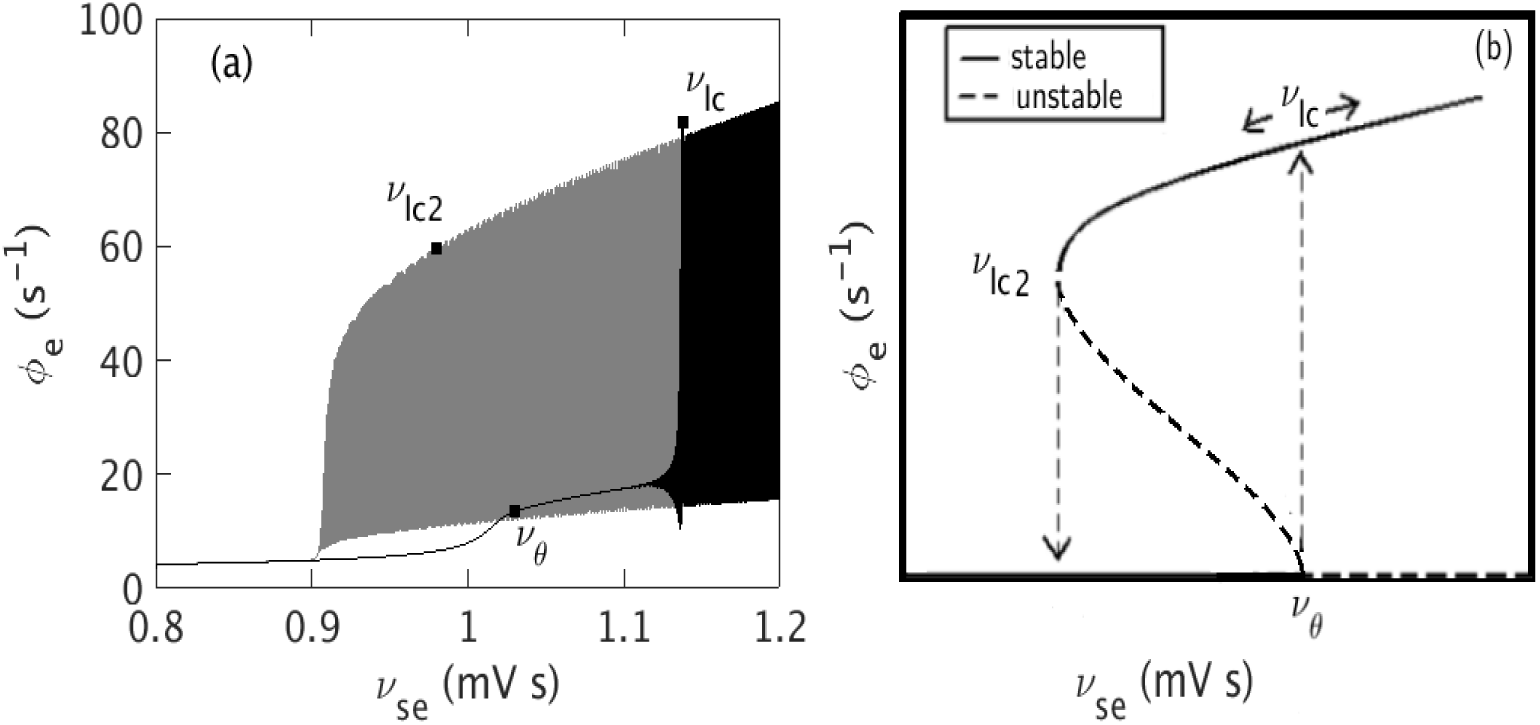
Hysteresis between seizure onset and offset. (a) *ν*_*se*_ vs. *ϕ*_*e*_. Black color shows the variation of *ϕ*_*e*_ during ramp up, i.e. during onset, and gray color shows the variation of *ϕ*_*e*_ during ramp down, i.e. during offset. (b) A schematic diagram of the hysteresis. Solid lines show stable states and dashed lines show unstable ones.

#### Analytical prediction of onset and offset transition times

Paralleling the analytic prediction of the characteristic time required to develop absence seizures [33], we next predict characteristic tonic-clonic onset and offset times.

For *ν*(*t*) ≈ *ν*_*θ*_, the oscillation amplitude *A* obeys

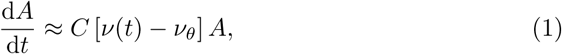

where *C* is a constant, and *ν*(*t*) is the instantaneous value of *ν*_*se*_. Because *ν*_*se*_ only varies with time *t*, we can make the approximation *ν*(*t*) − *ν*_*θ*_ ≈ *c*(*t* − *t*_*θ*_) near the threshold, when the oscillation starts at *A*_*θ*_. This yields

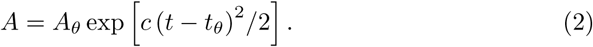

with 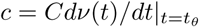 then *A* = *A*_*lc*_ at *t* = *t*_*lc*_

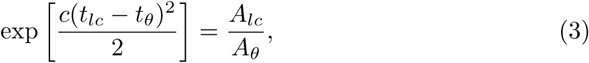

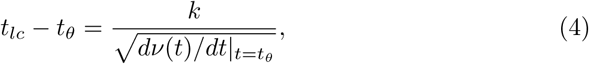

where *k* = [(2*/C*) ln(*A*_*lc*_*/A*_*θ*_)]^1/2^. Similar analysis predicts that the transition time *t*_*lc*_ − *t*_*sc*_ from the saddle-cycle attractor to the larger limit cycle also follows this scaling.

The decrease of oscillation amplitude during the ramp down period can be approximated as

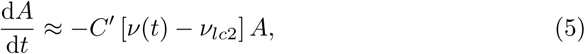

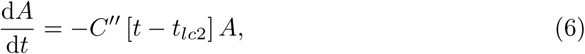

where *C*′ and *C*″ are constants, and *t*_*lc*2_ is the offset bifurcation threshold as mentioned in previous sections. This yields

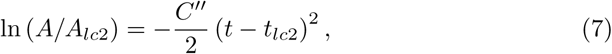

which indicates a superexponential decrease during seizure offset.

#### Dynamics during ictal state plateau

Figure 3 shows the phase space trajectory of *ϕ*_*e*_ for the default parameters in Table 1, except Δ = 2 s, which we use to see the saddle-cycle attractor more clearly. Figure 3(a) shows the trajectory of *ϕ*_*e*_ on the *ϕ*_*e*_ - d*ϕ*_*e*_/d*t* plane. In the left edge of the figure, we see the evolving fixed point, which first appears as straight line and then moves towards the right with increasing *ν*_*se*_. Once the system crosses the linear instability threshold, the fixed point becomes unstable and the trajectory spirals out to a large amplitude limit cycle attractor via an unstable saddle-cycle attractor. The amplitude of the large attractor increases gradually until *ν*_*se*_ = *ν*_max_, then decreases until *ν*_*lc*2_, where it becomes unstable and the system spirals back to the stable fixed point; no saddle-cycle is seen during the inward spiral. Three segments of the trajectory are shown in Figs 3(b) – (d), to clarify these dynamics. Figure 3(b) shows *ϕ*_*e*_ spiraling outward from the steady state to the saddle-cycle attractor with amplitude ≈ 30 s^−1^. Figure 3(c) shows the outward spiral from the saddle cycle to the limit cycle attractor with amplitude ≈ 90 s^−1^. Figure 3(d) shows the inward spiral during ramp down of *ν*_*se*_.

**Fig 3.**
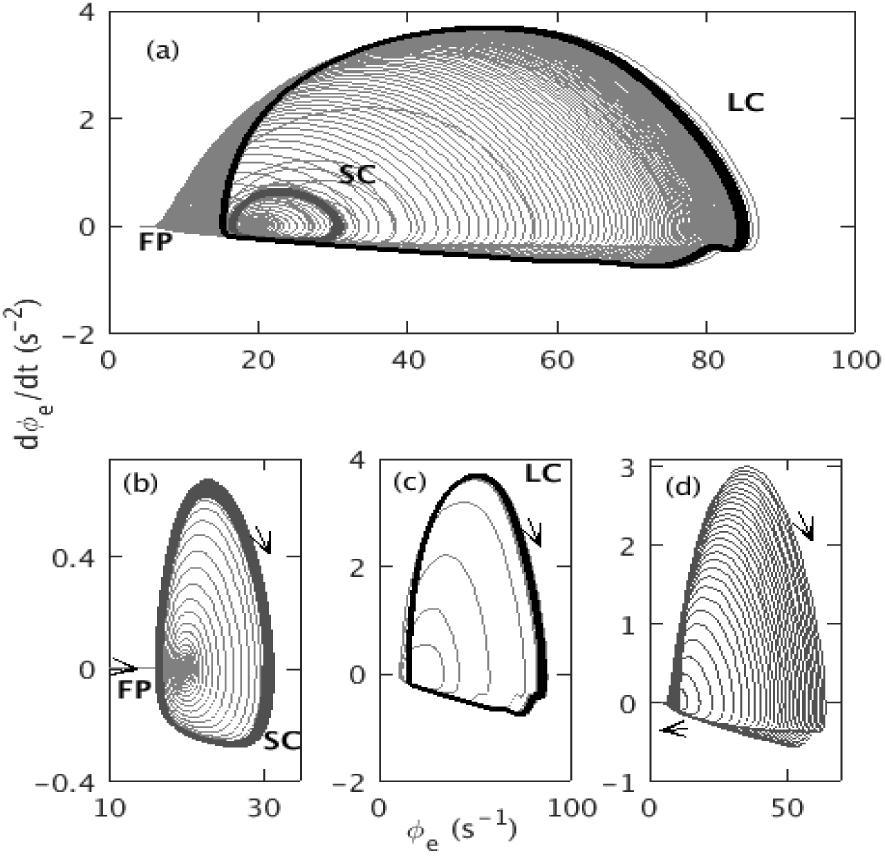
Phase space trajectory of *ϕ*_*e*_ for Δ = 2 s, and rest of the default parameters as in Table 1. (a) Trajectory from from *t* = 5 s to *t* = 295 s. Initial small straight line labeled with FP corresponds to the evolving fixed point; small dark gray segment labeled with SC corresponds to the saddle-cycle attractor; black segment labeled with LC corresponds to the large amplitude limit cycle attractor. The fixed point and center of the clockwise limit cycle trajectory move from left to right during ramp up and right to left during ramp down. (b) Trajectory from *t* = 104 s to *t* = 107 s. (c) Trajectory from *t* = 114.5 s to *t* = 150 s. (d) Trajectory from *t* = 200 s to *t* = 295 s.

Figure 4 shows the dynamic spectrum of *ϕ*_*e*_ from Fig. 1(b). A sudden appearance of 10 Hz oscillation with multiple harmonics at *t* = *t*_*θ*_ is seen. These harmonics resemble with the harmonics seen in [3], both experimentally and theoretically. The power of the harmonics decreases with harmonic number and their duration decreases slightly. We find a frequency broadening during the seizure onset at ∼ 113.5 s, due to the rapid change of the amplitude of the oscillations. Frequency broadening of the first few harmonics during seizure offset is also seen, and there is a slight frequency drop.

**Fig 4.**
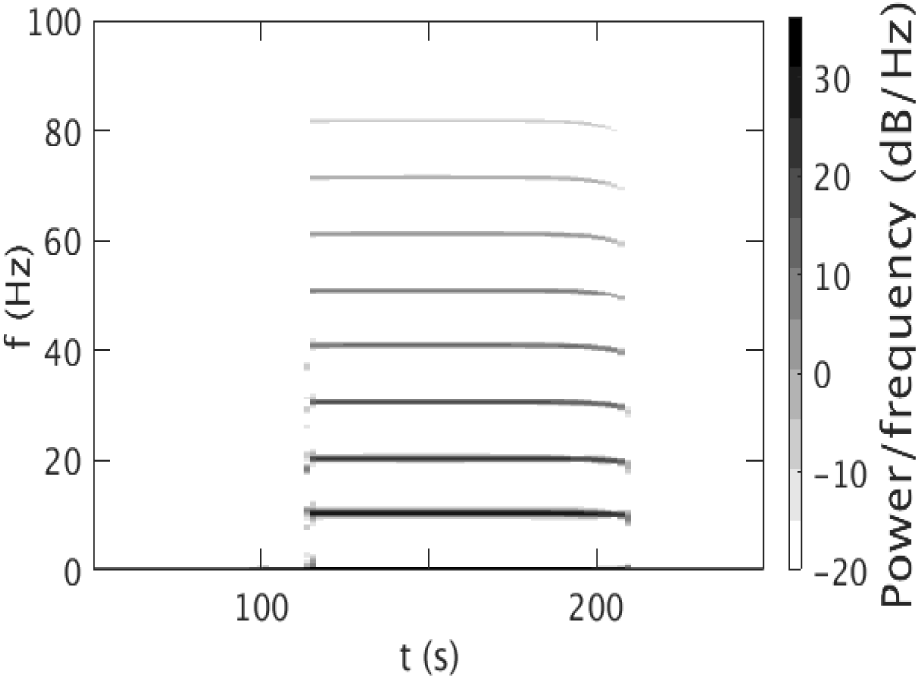
Dynamic spectrum for *ν*_max_ = 1.2 mV s with the parameters in Table 1. A Hanning window of 600 data points, an overlap of 200 points, and sampling frequency of 200 Hz was used. The color bar shows the dB scale.

#### Dynamics of corticothalamic seizure propagation

Figures 5 (a) and (b) show the time series of the fields *ϕ*_*r*_ during onset and offset, respectively. Similarly, Figs 5 (c) and (d) show the time series of the fields *ϕ*_*s*_ during onset and offset.

**Fig 5.**
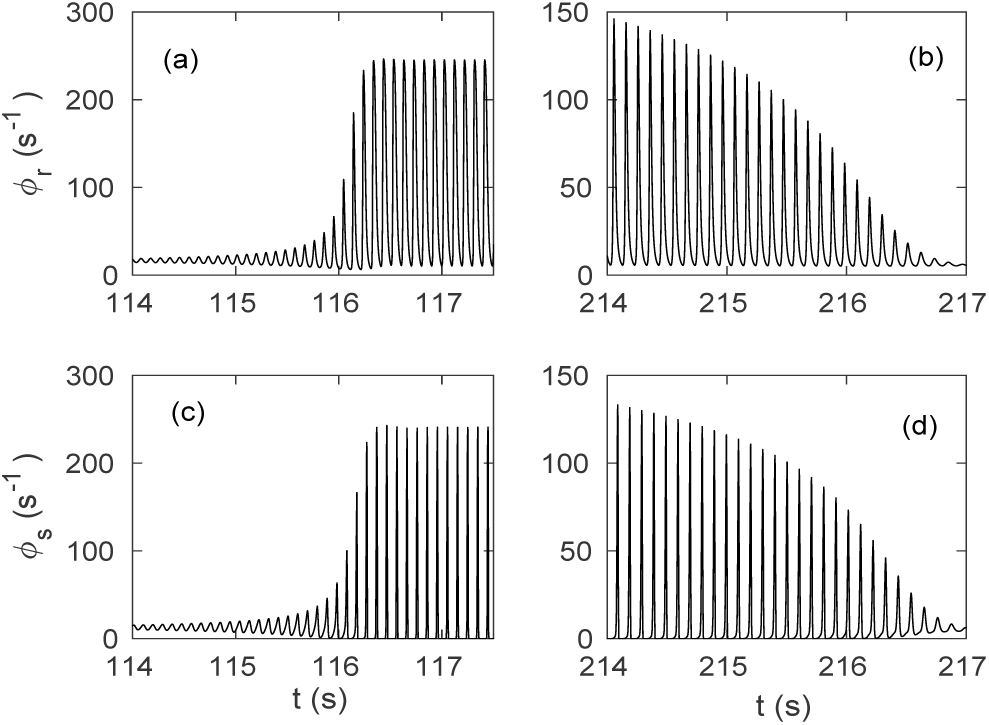
Time series of fields during seizure onset and offset: (a) *ϕ*_*r*_ at seizure onset. (b) *ϕ*_*r*_ at seizure offset. (c) *ϕ*_*s*_ at seizure onset. (d) *ϕ*_*s*_ at seizure offset.

From these plots we observe that (i) during onset *ϕ*_*r*_ reaches much higher amplitudes than *ϕ*_*e*_; and, (ii) the ratio between the amplitude of the small oscillations that develop after crossing the bifurcation and the amplitude of the saturated limit cycle is smaller for *ϕ*_*e*_ than it is for *ϕ*_*r*_ and *ϕ*_*s*_.

In order to study the interplay among *ϕ*_*e*_, *ϕ*_*r*_, and *ϕ*_*s*_ in more detail, we plot their limit cycle phase space trajectories and time series at *ν*_*se*_ ≈ *ν*_max_ in Fig. 6. Figures 6(a) and (b) show the time series and phase space trajectory of *ϕ*_*e*_, respectively. Figures 6(c) and (d) show the time series and phase space trajectory of *ϕ*_*r*_, respectively. A *t*_0_/2 time shift between the peaks of *ϕ*_*e*_ and *ϕ*_*r*_ is seen due to the propagation delay between these populations. We also see a wide minimum between two successive peaks of *ϕ*_*r*_. The phase space in Fig. 6(d) shows similar trajectory to Fig. 6(d), but with greater amplitude. Figures 6(e) and (f) show the time series and phase space of *ϕ*_*s*_, respectively, and they show an equal amplitude but wider peak than Figs 6(c) and (d). Figure 6 shows that all three fields exhibit slightly different trajectories, with the higher amplitudes of *ϕ*_*r*_ and *ϕ*_*s*_ near the maximum firing rate.

**Fig 6.**
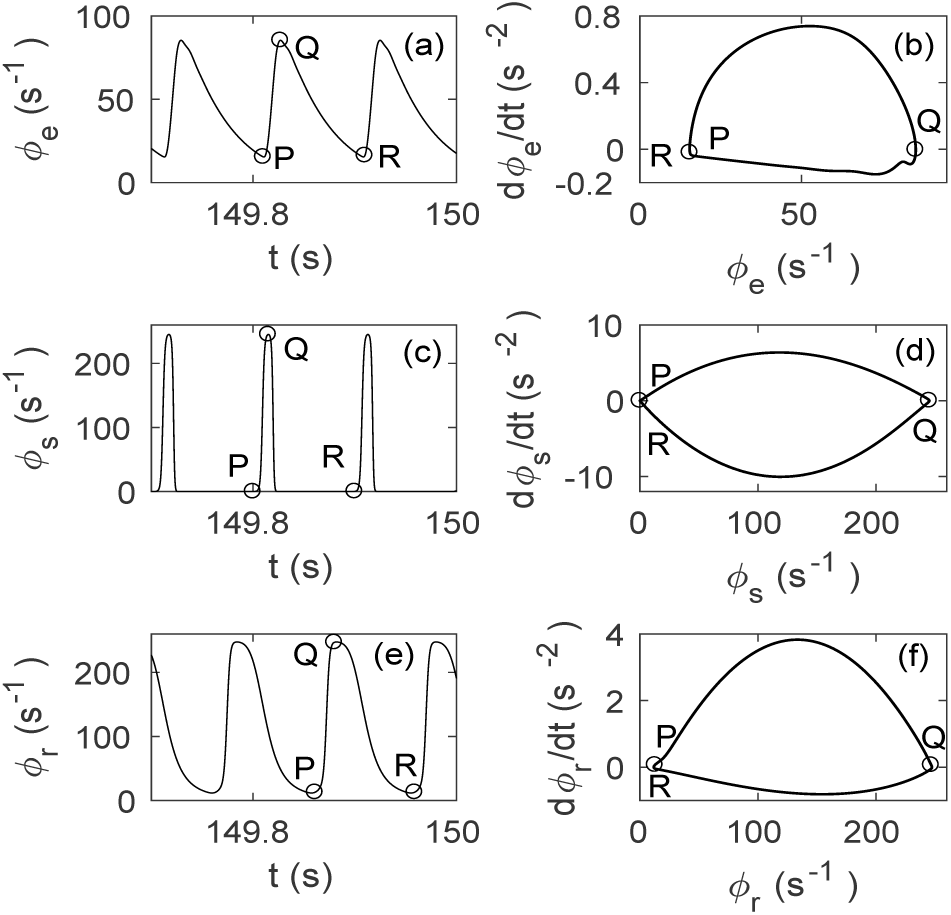
Mid-seizure limit cycle dynamics of *ϕ*_*e*_, *ϕ*_*s*_, and *ϕ*_*r*_ from *t* = 149.7 s to *t* = 150 s with other parameters as in Table 1. (a) Time series of *ϕ*_*e*_ at *ν*_*se*_ ≈ *ν*_max_. (b) Phase space trajectory of *ϕ*_*e*_. (c) *ϕ*_*r*_ at *ν*_*se*_ ≈ *ν*_max_. (d) Trajectory of *ϕ*_*r*_. (e) *ϕ*_*s*_ at *ν*_*se*_ ≈ *ν*_max_. (f) Trajectory of *ϕ*_*s*_. P and R are successive minimums and Q is the intermediate maximum.

Close examination of Fig. 6 reveals the signal flow through the populations. A peak of *ϕ*_*e*_ reaches *ϕ*_*r*_ and *ϕ*_*s*_ simultaneously *t*_0_/2 later. The peak of *ϕ*_*e*_ coincides approximately with the bottom of the trough of *ϕ*_*r*_, and a positive excitation with the maximum firing rate appears, which suppress *ϕ*_*s*_. This suppression then reduce the excitation of *ϕ*_*e*_ a time *t*_0_/2 later and causes an exponential decay. A negative perturbation to *ϕ*_*e*_ results, which then propagates to the thalamus again and reduces the excitation of *ϕ*_*r*_ after a further time *t*_0_/2, which allows a positive excitation of *ϕ*_*s*_ almost immediately. This positive excitation then flows to *ϕ*_*e*_ and initializes the next cycle of the loop.

In molecular level, the imbalance between inhibitory and excitatory conductances induced by blocking synaptic and voltage-gated inhibitory conductances, or by activating synaptic and voltage-gated excitatory conductances incorporates the positive feedback, which leads to seizures [17, 38]. Seizures are suppressed by the opposite manipulations: increasing inhibition or decreasing excitation [17, 38].

### Impact of temporal variation of *ν*_*se*_ on seizure dynamics

In this section, we investigate the effects of the temporal variation of *ν*_*se*_ on the model seizure dynamics by varying the maximum connection strength *ν*_max_, duration *t*_2_ − *t*_1_, and rise time Δ, holding all other parameters at the values in Table 1.

We first analyze the impact of the variation of *ν*_*se*_ on the overall dynamics of *ϕ*_*e*_, as shown in Fig. 7. For *ν*_max_ = 1 mV s in Fig. 7(a), *ϕ*_*e*_ increases with *ν*_*se*_ as shown in Fig. 16, then returns smoothly to the initial steady state value as *ν*_*se*_ returns to *ν*_0_. Figures 7(b) and (c) show that increasing *ν*_max_, yields periodic oscillations of increasing magnitude as corticothalamic feedback strengthens; oscillations also start earlier and are damped away later because the system crosses onset threshold earlier and offset threshold later for higher *ν*_max_. However, the system does not return to its initial steady state for *ν*_max_ *>* 1.542 mV s; instead it moves to the high firing steady state of Fig. 16.

**Fig 7.**
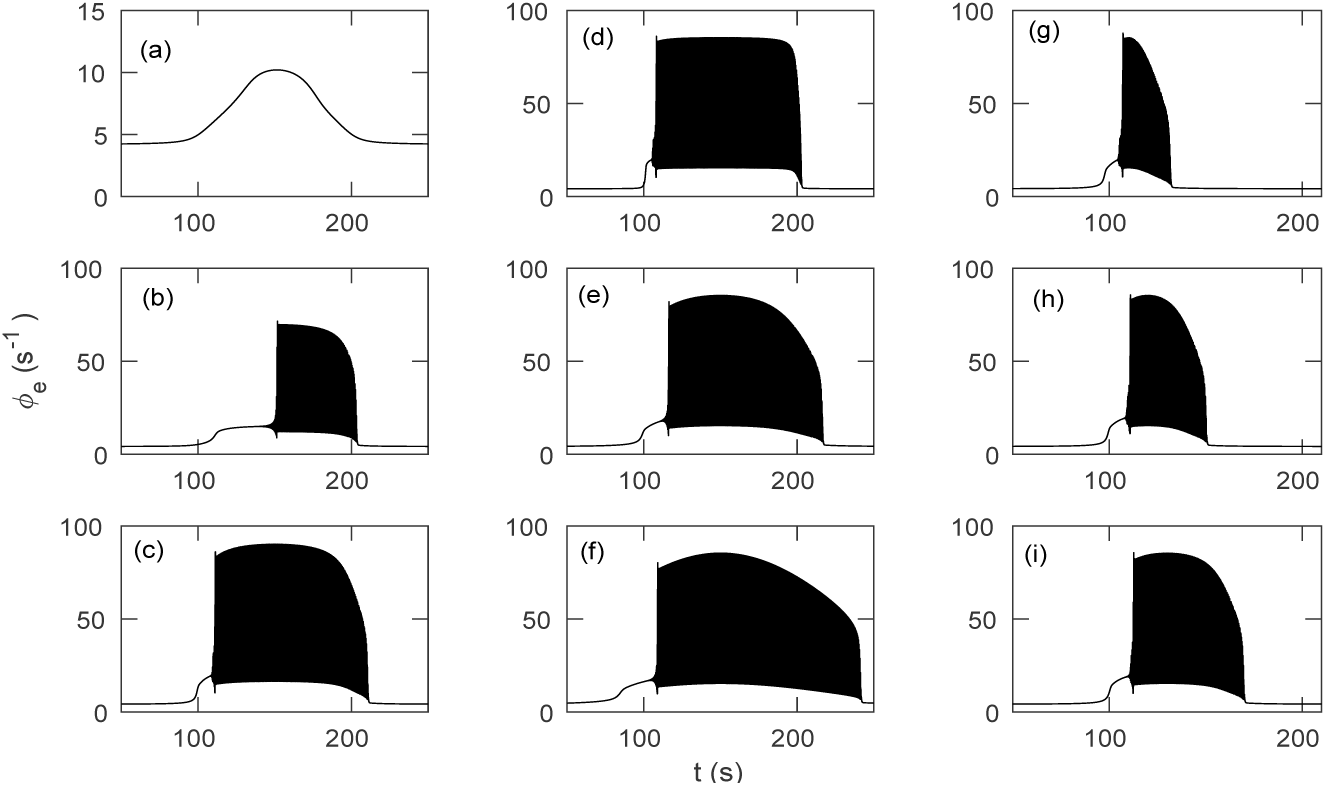
Time series for different temporal profiles of *ν*_*se*_, with other parameters as in Table 1. (a) *ϕ*_*e*_ vs. *t* for *ν*_max_ = 1 mV s. Individual oscillations cannot be distinguished. (b) *ν*_max_ = 1.05 mV s. (c) *ν*_max_ = 1.25 mV s. (d) Δ = 2 s. (e) Δ = 20 s. (f) Δ = 60 s. (g) *t*_2_ − *t*_1_ = 20 s. (h) *t*_2_ − *t*_1_ = 40 s. (i) *t*_2_ − *t*_1_ = 60 s.

Figures 7(d) – (f) show the effects of varying ramp width Δ from 2 s to 60 s. Figure 7(d) shows that for the step-like variation of *ν*_*se*_ for Δ = 2 s, the oscillations rapidly reach maximum amplitude after the transition to the large amplitude attractor and also decrease sharply from their maximum to the initial steady state once the system crosses the threshold during ramp down. Figures 7(e) and (f) show that the slower ramp for larger Δ implies that the amplitude of the oscillations during seizure onset and offset decreases more gradually.

Figures 7(g) – (i) show the effects of variation of the characteristic time *t*_2_ − *t*_1_ from 20 s to 100 s. As expected, the duration of seizure oscillations increases with *t*_2_ − *t*_1_.

#### Seizure onset time

Figure 8 quantifies the effects of *ν*_max_ and Δ on seizure onset. We do not revisit the variation with *t*_2_ − *t*_1_ because its effects were already discussed in the previous subsection.

**Fig 8.**
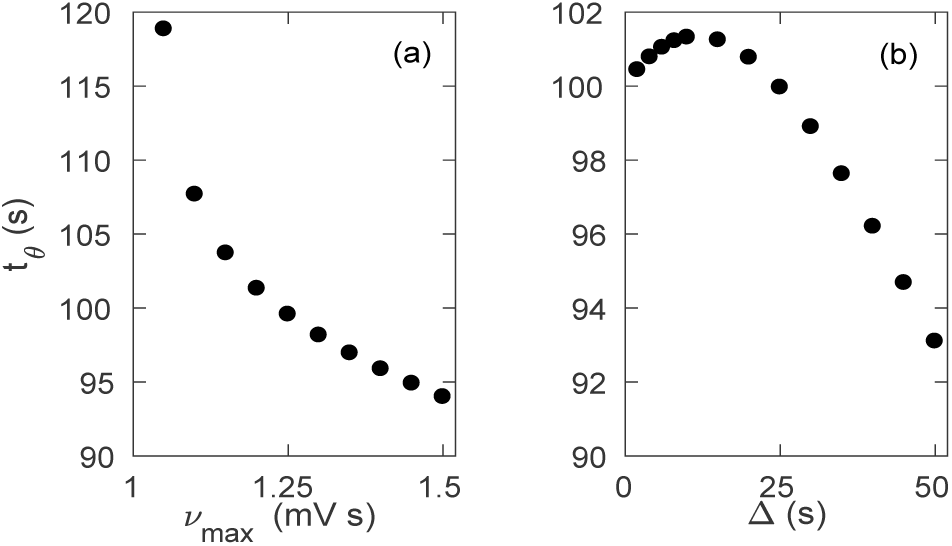
Effects of temporal variation of *ν*_*se*_ on seizure onset with parameters as in Table 1. (a) *t*_*θ*_ vs. *ν*_max_. (b) *t*_*θ*_ vs. Δ.

Figure 8(a) shows that *t*_*θ*_ decreases with increasing *ν*_max_, because the system reaches *ν*_*θ*_ earlier for a higher *ν*_max_. Figure 8(b) shows the variation of *t*_*θ*_ with Δ. For Δ *<* 10 s, *t*_*θ*_ increases slightly with Δ, because due to the high rate of change, *ν*_*se*_ rapidly approaches its maximum, crossing all the bifurcation values. At longer Δ *≥* 10 s, the temporal profile of *ν*_*se*_ becomes smooth and flat topped like Fig. 1(a) and *ν*_*se*_ gradually ramps up to the bifurcation point, so the system crosses the threshold later for a larger Δ, resulting in a decrease in *t*_*θ*_.

#### Dynamic spectrum

In this section we discuss the effects of changing the temporal profile of *ν*_*se*_ on the power spectrum of *ϕ*_*e*_ and use its evolution to further clarify the occurrence of saddle cycles.

Figure 9(a) shows the dynamic spectrum for *ν*_max_ = 1.05 mV s. During the seizure, we observe a peak at approximately ∼ 10 Hz with several harmonics. We also find lower frequency drop and broadening during seizure onset and offset as in Fig. 4. Figure 9(b) shows that for *ν*_max_ = 1.15 mV s, harmonics have greater duration and power than Fig. 9(a); frequency broadening is also more prominent. Figure 9(c) shows that for *ν*_max_ = 1.55 mV s, there is no oscillation after *t* = 143.52 s. A detailed investigation shows that the power of the peaks increases significantly with *ν*_max_ and *t*_2_ − *t*_1_, but decreases slightly with Δ, especially at higher order harmonics. A small peak around 205 s shows that the system returns to the initial steady state via small oscillation after it crosses the offset bifurcation.

**Fig 9.**
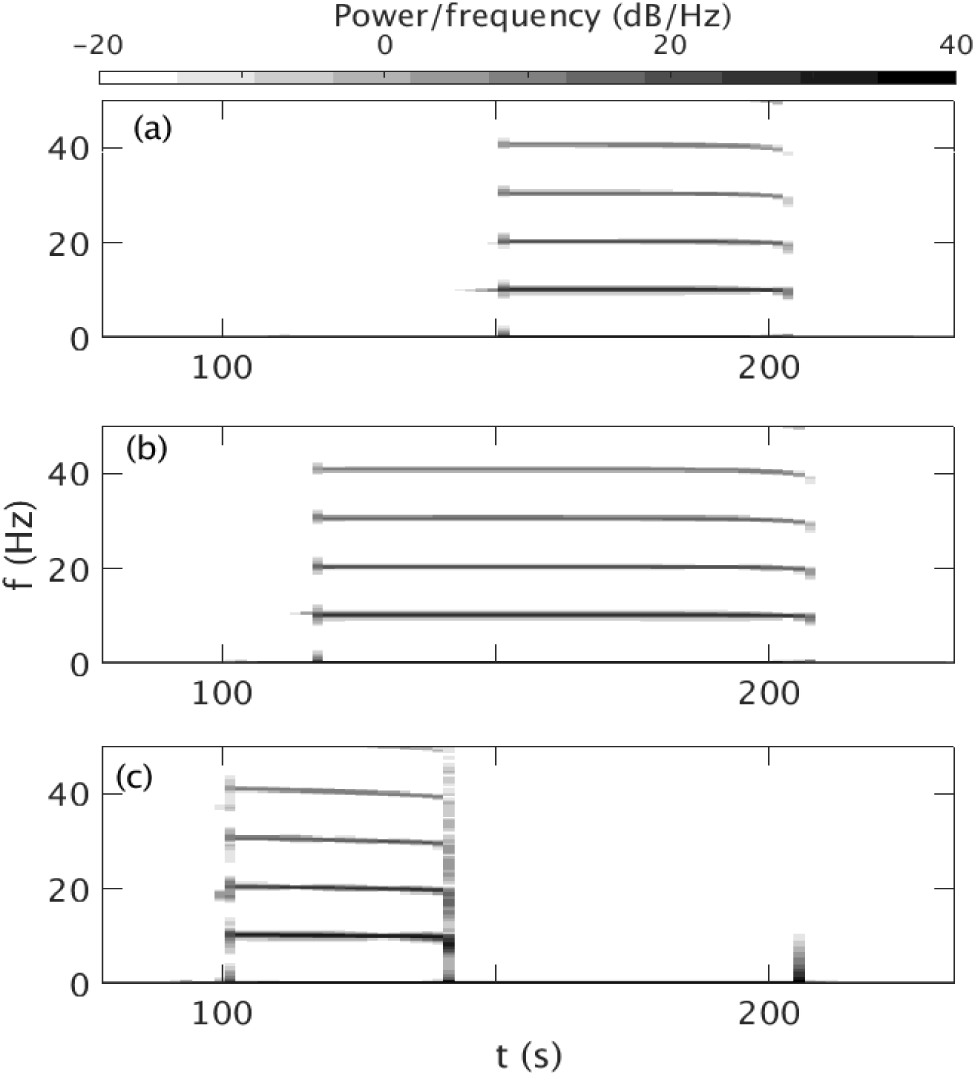
Dynamic spectrum vs. *ν*_max_ for the parameters in Table 1. The power density of the harmonics is calculated using a Hanning window of 600 data points, an overlap of 200 points, and sampling frequency of 200 Hz, the color bar at top shows the dB scale. (a) Dynamic spectrum for *ν*_max_ = 1.05 mV s. (b) *ν*_max_ = 1.15 mV s. (c) *ν*_max_ = 1.55 mV s.

#### Characteristic transition times

In this section we test the analytic prediction made in earlier sections. Figure 10(a) shows *t*_*lc*_ − *t*_*θ*_ vs. (*dν*_*se*_*/dt*)^−1/2^. A least-squares fit to these data yields

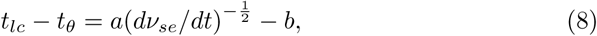

with *a* = (0.042 ± 0.004) V^1/2^ s and *b* = (0.9 ± 1.4) s, which is consistent with Eq. (4). Figure 10(b) shows (*dν*_*se*_*/dt*)^−1/2^ vs. *t*_*lc*_ − *t*_*sc*_. A least-squares fit yields

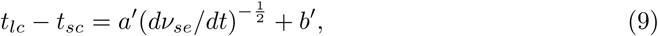

with *a*′ = (0.003 ± 0.001) V^1/2^ s and *b*′ = (0.0 ± 0.2) s, which has the same scaling as Eq. (4).

**Fig 10.**
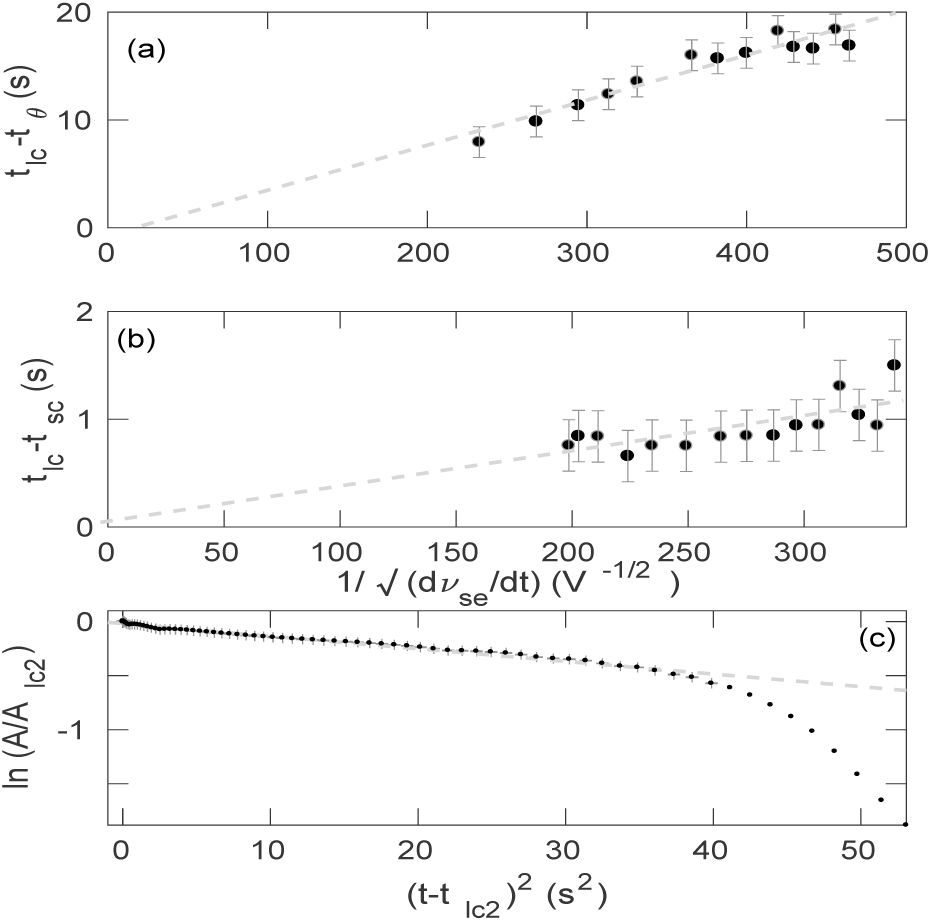
Dependence of seizure transition times on (*dν*_*se*_*/dt*)^−1/2^ with the default parameters as in Table 1 and Δ ranges from 2 s to 50 s. (a) *t*_*lc*_ − *t*_*θ*_ vs. (*dν*_*se*_*/dt*)^−1/2^; (b) *t*_*lc*_ *t*_*sc*_ vs. (*dν*_*se*_*/dt*)^−1/2^, and (c) ln(*A/A*_*lc*2_) vs. (*t t*_*lc*2_)^2^ for Δ = 10 s and time ranges from 190 s to 250 s. Error bar represent uncertainties of the least-squares fits. Points with no error bars are not considered for the least-squares fit.

Figure 10(c) shows ln(*A/A*_*lc*2_) vs. (*t* − *t*_*lc*2_)^2^ for Δ = 10 s, which follows Eq. (7) until the amplitudes of the oscillations start to decrease super-exponentially towards the steady state. A least-squares fit to the linear decrease yields

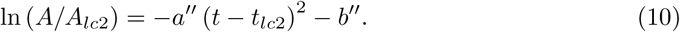

with *a*″ = (0.0116 ± 0.0002) s^−2^ and *b*″ = (0.018 ± 0.004). The figure shows that the decrease of the envelope follow the linear fit for a relatively short time, after which the decrease becomes steeper. By using Eqs (2) and (3), it can be also shown that decrease within the linear region also follows the same scaling as Eq. (4).

#### Saddle Cycle

Previously, we mentioned the presence of a small amplitude ∼ 18 Hz saddle cycle. The system orbits there for few seconds, then spirals out towards the large amplitude limit cycle attractor. However, this saddle-cycle is not observed in all cases, for example, a colose zoom near the onset of all subfigures of Fig. 7 will show that the small amplitude saddle-cycle oscillations like Fig. 1(c) are only prominent in Figs 7(c) and (d). Here, we explore the dependence of the saddle-cycle oscillations on *ν*_max_ and Δ.

Figure 11 shows the variation of saddle-cycle oscillations with respect to *ν*_max_, with other parameters as in Table 1. Figure 11(a) shows the phase space trajectory for *ν*_max_ = 1.15 mV s. No saddle-cycle attractor is seen in this figure. Figure 11(b) shows the trajectory for *ν*_max_ = 1.25 mV s. A small saddle-cycle attractor is seen between the fixed point and the large amplitude attractor. Figures 11(c) and (d) show the trajectories for *ν*_max_ = 1.35 mV s and 1.45 mV s, respectively. The saddle cycle increases in size with *ν*_max_. A similar investigation shows that similar phenomena occur when Δ is varied, with the saddle cycle being most prominent for small Δ, completely disappearing for Δ ≳ 20 s.

**Fig 11.**
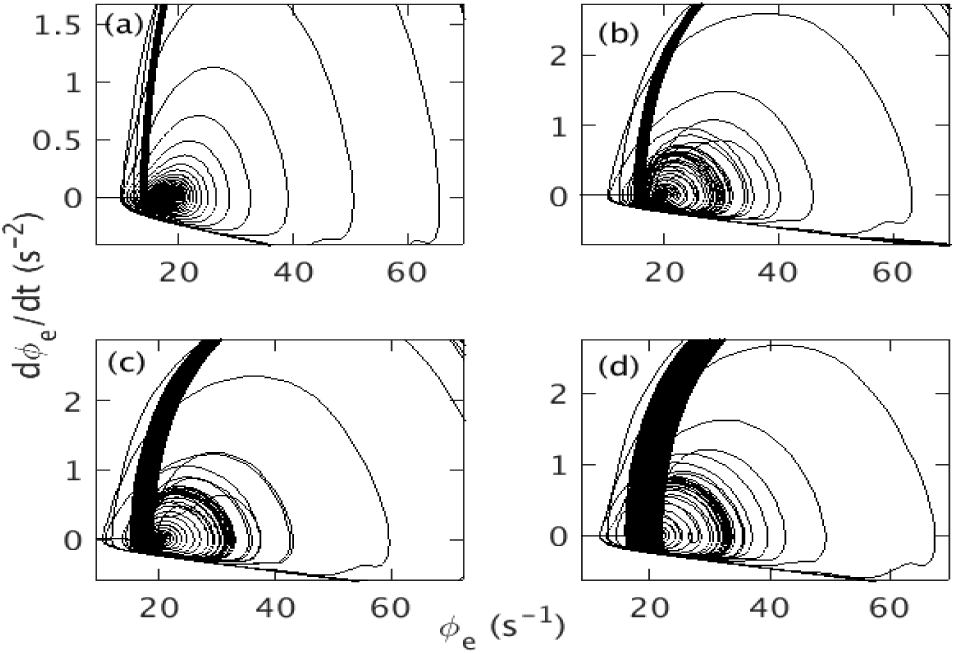
Effects of variation of *ν*_max_ on saddle-cycle with rest of the parameters as in Table 1. (a) Phase space trajectory for *ν*_max_ = 1.15 mV s. (b) Trajectory for *ν*_max_ = 1.25 mV s.(c) Trajectory for *ν*_max_ = 1.35 mV s. (d) Trajectory for *ν*_max_ = 1.45 mV s.

To understand the relation between the saddle-cycle oscillation and rate of change of *ν*_*se*_ more clearly, we calculate the power spectrum for different *ν*_max_ and Δ. Figure 12(a) shows the variation of the power spectrum with *ν*_max_. For a small *ν*_max_, there is no peak around 18 Hz, but a peak at approximately 18 Hz appears when *ν*_max_ *≥* 1.2 mV s and becomes more prominent and strong with increasing *ν*_max_. Figure 12(b) shows that the power of the peak around 18 Hz decreases with Δ and disappears for Δ ≳ 20 s.

**Fig 12.**
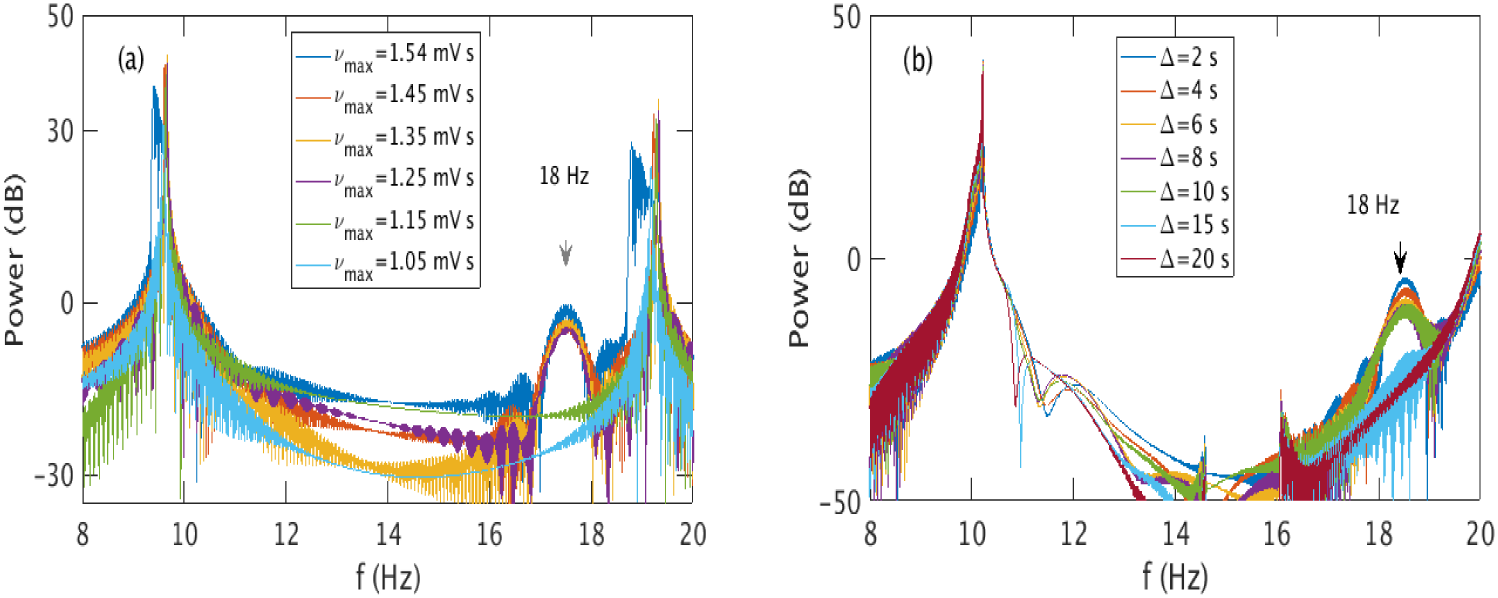
(Color online) Variation in the power of the saddle-cycle oscillations with rest of the parameters in Table 1. (a) Power spectrum vs. *ν*_max_. (b) Power spectrum vs. Δ. Legends show the corresponding values of *ν*_max_ and Δ.

These results imply that the presence of saddle-cycle oscillations depends on the rate of change of of *ν*_*se*_. Figure 13 illustrates the presence or absence of saddle-cycle oscillations for 236 different combinations of *ν*_*se*_ and Δ as a function of the value of *dν*_*se*_*/dt*. When *dν*_*se*_*/dt <* 7 *×* 10^−3^ mV, there are no saddle-cycle oscillations; for *dν*_*se*_*/dt >* 9 *×* 10^−3^ mV, the system always exhibits saddle-cycle oscillations; while for 7 *×* 10^−3^ ≲ *dν*_*se*_*/dt* ≲ 9 *×* 10^−3^ mV, there is a narrow mixed region where the presence of saddle cycle cannot be predicted solely from the rate of change of *ν*_*se*_.

**Fig 13.**
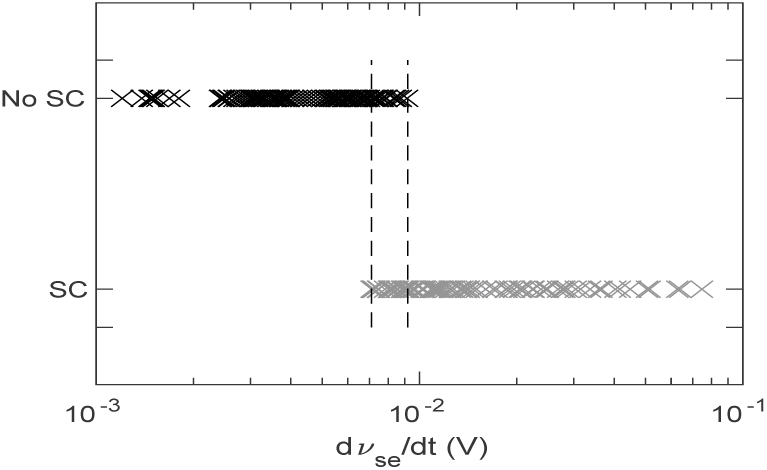
Dependence of saddle-cycle oscillations on *dν*_*se*_*/dt*. Gray crosses show the presence of a saddle-cycle and black crosses show its absence.

In order to see why saddle cycles are only seen for high *dν*_*se*_*/dt*, we show the time evolution of 10 Hz and 18 Hz frequency peaks for Δ = 2 s and Δ = 50 s in Fig. 14 during seizure onset with other parameters as in Table 1. In Fig. 14(a), for Δ = 50 s and *dν*_*se*_*/dt* = 0.003 mV, the 10 Hz peak always rise faster than the 18 Hz peak, and hence, always has more power and dominates the spectrum; no saddle cycles are seen in the trajectory. On the other hand, in Fig. 14(b), for Δ = 2 s and *dν*_*se*_*/dt* = 0.03 mV, the 18 Hz peak rises faster than the 10 Hz peak during onset so there is a ∼ 2 s window in which the 18 Hz peak dominates and hence, the system is seen to exhibit saddle-cycle oscillations during onset in Fig. 1, after which the 10 Hz peak dominates. Now, since, *ν*_*θ*_ is a the bifurcation threshold and does not depend on the temporal profile, but *ν*_*lc*_ depends on the temporal profile and the time to reach the 10 Hz limit cycle (i.e., *t*_*lc*_ − *t*_*θ*_), we conclude that *ν*_*lc*_ is the parameter that defines the existence of the saddle cycle. The system will exhibit saddle cycle oscillation only if *ν*_*sc*_ *> ν*_*lc*_ at *t*_*sc*_.

**Fig 14.**
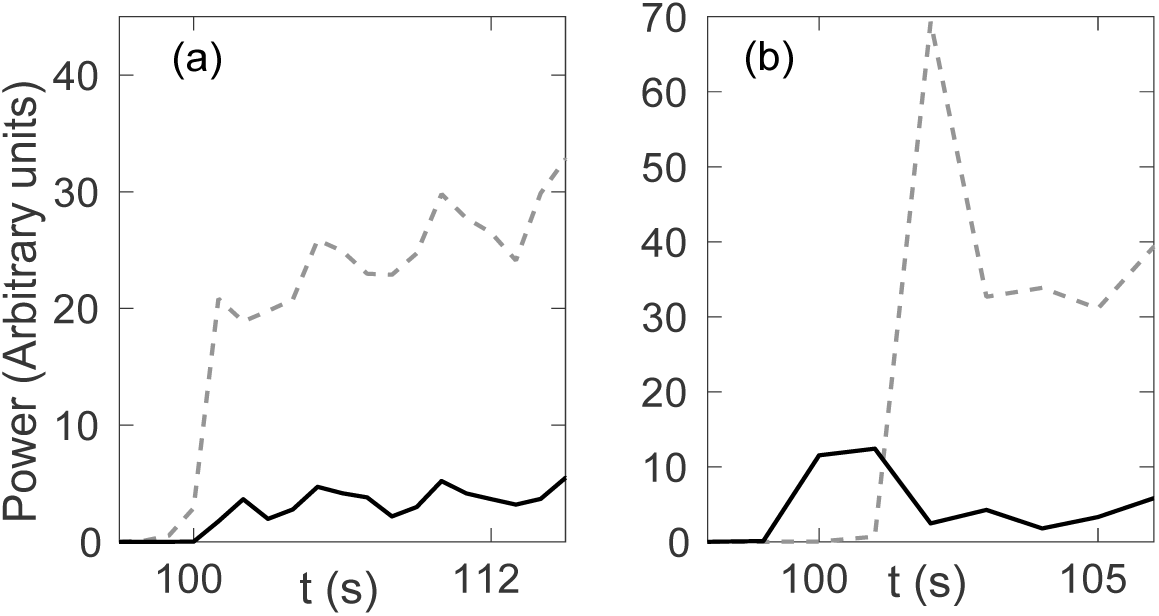
Temporal variation of frequency peaks during seizure onset; black solid line shows the ∼ 18 Hz peak; gray dashed line shows the ∼ 10 Hz peak with parameters from Table 1. (a) Δ = 50 s; (b) Δ = 2 s.

## Discussion

We have used an established neural field model of the corticothalamic system [3] to study the dependence of tonic-clonic seizures on the temporal profile of a corticothalamic connection strength *ν*_*se*_ that induces seizures. The effects of varying other connection strengths can also be qualitatively predicted using these outcomes because they will exhibit similar dynamics due to the universality properties of the Hopf bifurcation. Also, the function [Eq. (20)] used to vary the connection strength is an approximation of what seems to occur in living systems. This function is an improvement over previous piece-wise linear functions [3]. The parameters and the shape of Eq. (20) could be customized in the future using experimental data. The key outcomes are:

i. The system exhibits ∼10 Hz limit cycle oscillations once the connection strength crosses the bifurcation threshold of *ν*_*θ*_ = 1.025 mV s, which is the characteristic frequency of tonic-clonic seizure via a subcritical Hopf bifurcation. The system returns to the resting equilibrium when the connection strength decreases below the offset threshold, *ν*_*lc*2_ = 0.98 mV s. The difference in onset and offset bifurcation values causes hysteresis; consistent with previously published results that used piecewise linear variation of *ν*_*se*_, rather than the present more realistic continuous gradual variation.
ii. For *ν*_max_ ≳ 1.542 mV, the system moves to another steady state near maximum firing rate and only returns to the initial steady state once *ν*_*se*_ returns below an offset threshold.
iii. The amplitude of *ϕ*_*e*_ increases with the maximum connection strength, *ν*_max_, because an increase of the connectivity strength increases the strength of the positive feedback loop between the cortex and the thalamus.
iv. Because increasing the maximum connection strength *ν*_max_ increases the amplitudes of the oscillations, it increases the power and the characteristic number of harmonics. The power of the harmonics also increases with the seizure duration *t*_2_ − *t*_1_, but decreases slightly with the ramp duration Δ.
v. The characteristic transition times required to reach the saturated limit cycle oscillation from the seizure threshold or the end of the saddle-cycle oscillations to the steady state are predicted and verified numerically to be inversely proportional to the square root of the rate of change of the connection strength.
vi. The system can also show transient ∼ 18 Hz saddle-cycle oscillation at the beginning of the seizure for high *dν*_*se*_*/dt* before moving to the 10 Hz attractor. These saddle-cycles become more prominent as *dν*_*se*_*/dt* increases; a system with *dν*_*se*_*/dt <* 7 *×* 10^−3^ mV never exhibits saddle-cycles, whereas one with *dν*_*se*_*/dt >* 9 *×* 10^−3^ mV always does.

Overall, the present study enables the varying spectral and temporal characteristics of seizures to be related to underlying physiological changes of the brain, such as changes in the connection strength between the cortex and the thalamus. The outcomes can be used for explaining the variability of seizure onset properties and seizure frequency across subjects by examining the temporal and spectral characteristics of seizure [39–41]. It may thus be possible to constrain the physiological properties of the corticothalamic connection strength dynamics of a subject by comparing the wave properties of seizure oscillations, such as amplitude, and frequency, with theory. Real-time fitting of the theoretical dynamics to observed waveforms may also be feasible, leading to the possibility of implementing feedback control systems based on the dynamics.

## Methods

In this section, we present a brief description of the corticothalamic neural field model used, along with the form of temporal variation of corticothalamic coupling strength [3, 4, 8].

### Corticothalamic Field Model

To investigate the dynamics of tonic-clonic seizure, we use the neural field model of the corticothalamic system seen in Fig. 15. In this study we use the same analytical model of [33], but in different parametric regime suitable to study the tonic-clonic seizure. The neural populations are denoted as: *e* = excitatory cortical; *i* = inhibitory cortical; *s* = thalamic relay neurons; *r* = thalamic reticular nucleus; and *n* = external inputs. The dynamical variables within each neural population *a* are the local mean cell-body potential *V*_*a*_, the mean rate of firing at the cell-body *Q*_*a*_, and the propagating axonal fields *ϕ*_*a*_. The firing rates *Q*_*a*_ are related to the potentials *V*_*a*_ by the response function

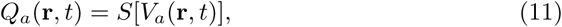

where *S* is a smooth sigmoidal function that increases from 0 to *Q*_max_ as *V*_*a*_ increases from −∞ to ∞, with

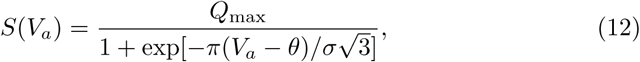

where *θ* is the mean neural firing threshold, *σ* is the standard deviation of this threshold, and *Q*_max_ is the maximum firing rate [3, 8].

In each neural population, firing rates *Q*_*a*_ generate propagating axonal fields *ϕ*_*a*_ that approximately obey the damped wave equation [3, 8]

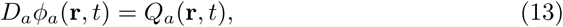

where the spatiotemporal differential operator *D*_*a*_ is

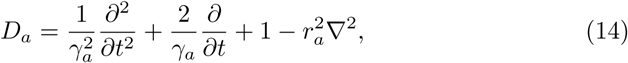

where *γ*_*a*_ = *v*_*a*_*/r*_*a*_ is the damping rate, *r*_*a*_ and *v*_*a*_ are the characteristic range and conduction velocity of axons of type *a*, and ∇^2^ is the Laplacian operator. The smallness of *r*_*i*_, *r*_*s*_, and *r*_*r*_ enables us to set *γ*_*a*_ ≃ ∞ except for *a* = *e*. The cell-body potential *V*_*a*_ results after postsynaptic potentials have propagated through the dendritic tree and then been summed as their resulting currents charge the soma. For excitatory and inhibitory neurons within the cortex, this is approximated via the second-order delay-differential equation [8]

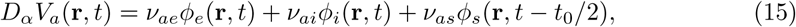

where *a* = *e, i* and the temporal differential operator is given by

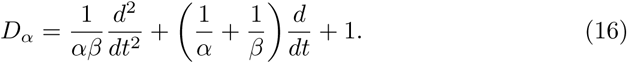

**Fig 15.**
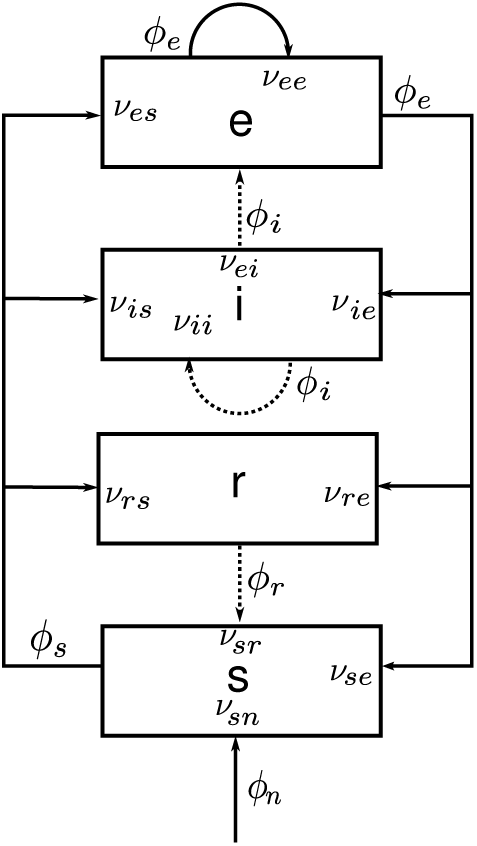
Schematic diagram of the corticothalamic model system. The neural populations shown are cortical excitatory (*e*), inhibitory (*i*), thalamic reticular (*r*), thalamic relay (*s*), and *n* = external inputs. The parameter *ν*_*ab*_ quantifies the connection to population *a* from population *b*. Inhibitory connections are shown with dashed lines.

The quantities *α* and *β* in Eq. (16) are the inverse decay and rise times, respectively, of the cell-body potential produced by an impulse at a dendritic synapse. Note that input from the thalamus to the cortex is delayed in Eq. (15) by a propagation time *t*_0_/2. For neurons within the specific and reticular nuclei of the thalamus, it is the input from the cortex that is time delayed, so

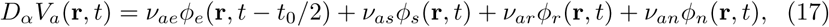

where *a* = *s, r*. The connection strengths are given by *ν*_*ab*_ = *N*_*ab*_*s*_*ab*_, where *N*_*ab*_ is the mean number of synapses to neurons of type *a* from type *b* and *s*_*ab*_ is the strength of the response in neurons *a* to a unit signal from neurons of type *b*. The final term on the right-hand side of Eq. (17) describes inputs from outside the corticothalamic system. In order to simplify the model we only include the connections shown in Fig. 15, so only 10 of the possible 16 connections between the four neural populations are nonzero [8]. We also assume the random intracortical connectivity and the number of connections between populations is proportional to the number of synapses [42, 43]. This random connectivity assumption provides *N*_*ib*_ = *N*_*eb*_ for all *b*, so *ν*_*ee*_ = *ν*_*ie*_, *ν*_*ei*_ = *ν*_*ii*_ and *ν*_*es*_ = *ν*_*is*_ [30].

Setting all spatial and temporal derivatives in Eqs (12) − (17) to zero determines spatially uniform corticothalamic steady states. The steady state firing rate, 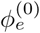 of *ϕ*_*e*_ is then given by [18]

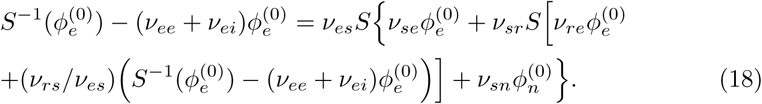

The properties of steady states in the corticothalamic model have been studied extensively in [8, 18], and we use the outcomes to identify the stable and unstable regions of the steady state. Figure 16 shows the steady state dependence of 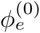 on *ν*_*se*_ with other parameters as in Table 1. It is seen that there are two stable steady state solutions: one corresponds to low mean firing rate and another to very high mean firing rate [18]. The low firing steady state was identified with normal states of brain activity in previous studies [8, 26]. The low firing-rate fixed point loses its stability at *ν*_*se*_ = *ν*_*θ*_. A steep increase in 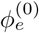 is seen near *ν*_*i*_ because the increasing *ν*_*se*_ push the sigmoid from its minimum by increasing the 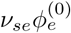 in Eq. (18), which results in an increase of the gain between the thalamus and the cortex. With further increase of *ν*_*se*_, the system eventually moves to a steady state with near-maximum firing rate. This high firing steady state is beyond the scope of our model because it will lead to effects such as hypoxia, which are not included here.

**Fig 16.**
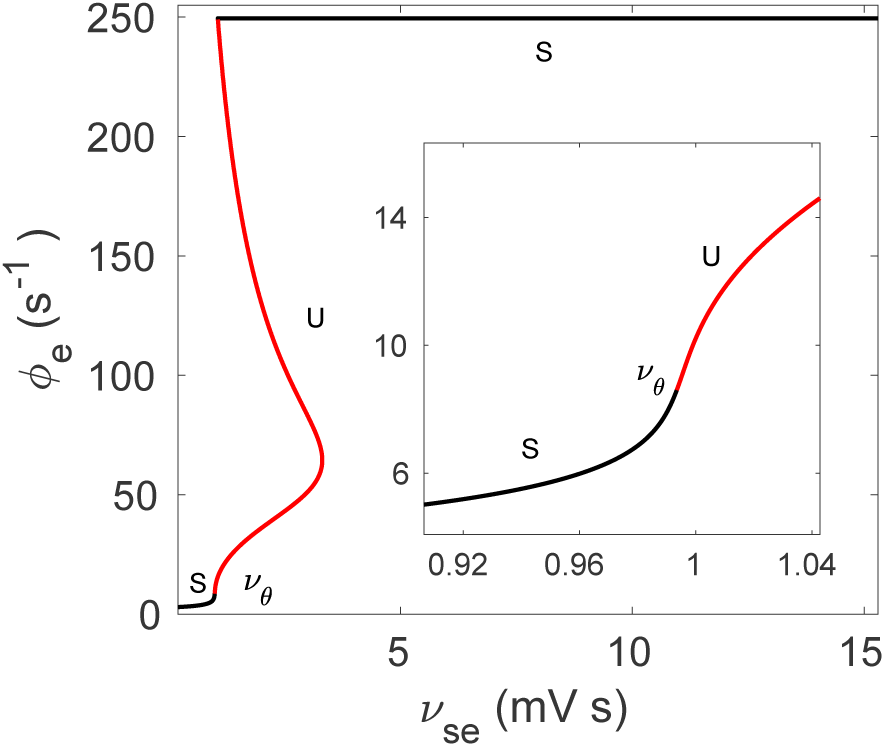
(color online) Steady states solution of the corticothalamic system for the variation of *ν*_*se*_ for tonic-clonic seizure. Black lines and the letter ‘S’ represent the stable steady state, and red lines and the letter ‘U’ represent the unstable steady states. Here *ν*_*θ*_ is the threshold value when the stable steady state becomes unstable. The inset shows zoomed view of the area around *ν*_*θ*_.

### Temporal Ramping

Brain activity propagates via the coupling of the various neuronal populations. Previous studies have shown that a gradual ramp-up of the coupling strength between the neuronal populations can lead from a stable steady state to periodic seizure oscillations [3, 33]. It is also seen that the dynamical and spectral characteristics of the resultant seizure-like oscillations depend on the physiological properties of the ramp of the coupling strength, such as, the maximum amplitude of the ramp, ramp rate, and characteristic duration [33].

In this paper, we ramp the coupling strength *ν*_*se*_ from an initial value *ν*_0_ to a maximum value *ν*_max_ and back to see the impact of the ramp characteristics on tonic-clonic seizures, with [33]

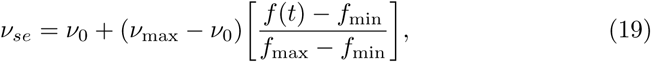

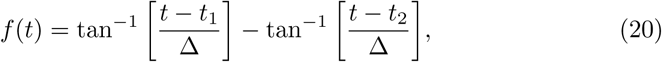

where *t* is the time. The ramp rise is centered on *t*_1_, and the ramp fall is centered on *t*_2_, and Δ is the characteristic rise time. Now, 0 ⩽ *f* (*t*) ⩽ *π*, so we normalize by dividing by *f*_max_ − *f*_min_ as seen in Eq. (19), where *f*_max_ and *f*_min_ are the maximum and minimum values of *f* (*t*) actually encountered in a given instance.

### Numerical Methods

We use *NFTsim* [44] to solve Eqs (11) – (17) numerically for the spatially uniform case in which the ∇^2^ term in Eq. (14) is zero. To vary *ν*_*se*_ temporally, we use Eqs (19) and (20). This involves solving ordinary delay differential equations, because there is a propagation time delay *t*_0_/2 between the different neural populations present in Eqs (15) and (17). Hence, a fourth-order Runge-Kutta integration is employed to solve these equations, with an integration time step of 10^−4^ s and store time histories of the delay terms *t*_0_/2 into the past.

Because extensive comparisons with experiment have demonstrated that the normal brain operates close to stable fixed points [3, 8, 18, 30, 32], we start our simulations from a corticothalamic steady state with low firing rate. However, because of the delay time *t*_0_/2, we must specify these initial steady-state conditions to apply for times −*t*_0_/2 < *t* ⩽ 0.

We use the parameters in Table 1 as the initial parameters, which are taken from [3] with *ν*_0_ = 0.8 mV s in all cases. A constant input *ν*_*sn*_*ϕ*_*n*_ = 2 mV is used and no external noise is applied in the simulations as the seizure onset occurs spontaneously.

Simulations are 300 s long, and we record the output time series every 5 ms. For all simulations, we use the default parameters shown in Table 1 unless otherwise specified. The default parameters we used are the corresponding parameter set of [3] for tonic-clonic seizure which push the system into the vicinity of alpha instability. For the dynamic spectrum and power spectrum analysis, we employ the FFT (fast Fourier transform) algorithm with a Hanning window of 600 data points with an overlap of 200 points and sampling frequency of 200 Hz.

## Acknowledgments

This work was supported by the Australian Research Council Center of Excellence for Integrative Brain Function (ARC Center of Excellence Grant CE140100007) and by an Australian Research Council Laureate Fellowship (Grant FL140100025).

